# An individual, mechanistic and dynamical model to simulate urban tree growth and ecosystem services supply under future scenarios

**DOI:** 10.1101/2025.10.09.681179

**Authors:** Davide Stucchi, Javier Babí Almenar, Renato Casagrandi

## Abstract

Urban trees represent a key nature-based solution and an essential component of green infrastructures, providing multiple ecosystem services but increasingly subjected to environmental stressors. Here we present a dynamic, mechanistic, and individual-based model designed to simulate growth of urban trees and the associated provision of ecosystem services under varying climate conditions. The model is modular, runs at daily time steps and incorporates key biological processes such as photosynthesis, water limitation and biomass allocation. It simulates single-tree growth and quntifies ecosystem services, such as carbon sequestration, air filtration, and local climate regulation using species-specific parameters and local climate forcing. The model was calibrated and tested in a pilot application in Milan, simulating the long-term growth of three of three common broadleaved species (*Platanus x acerifolia, Populus nigra* and *Robinia pseudoacacia)* across different planting ages and climate scenarios. A multi-objective calibration was used to fit stem diameter and crown width, and sensitivity analyses were conducted to assess parameter robustness and uncertainty propagation. Results show realistic growth trajectories with clear species–age contrasts. Tree growth declines under stronger climate forcing, and ecosystem service provision scales non-linearly with age, with mature trees delivering far greater benefits. Carbon sequestration and air filtration decrease under more extreme scenarios, whereas local climate regulation exhibits a compensatory response: lower productivity is offset by higher evaporative demand, yielding stable or slightly increased evapotranspiration-based cooling. The model offers a promising tool for supporting urban forestry decisions related to planning, species selection, and long-term ecosystem service provision.

## 1. Introduction

Urban areas are increasingly exposed to environmental challenges driven by climate change, including intensified heat waves, altered precipitation patterns, and degraded air quality (Cheval et al., 2024; Intergovernmental Panel on Climate Change, 2023). In this context, urban greening, has emerged as a critical action to locally mitigate some of these impacts, by regulating microclimate, enhancing air filtration, and improving stormwater retention (Babí Almenar et al., 2021; Ernst et al., 2022; Pugh et al., 2012). These positive outcomes are often referred to as ecosystem services (ES). Among the different components of urban greening, trees play a central role: due to their size, structure, and longevity, they provide disproportionate levels of ES compared to other vegetated features, and act as long-term regulators of urban environmental quality (Pataki et al., 2021; Pearlmutter et al., 2017). However, in order to quantify and plan for these services, especially under uncertain future conditions, it is essential to model the dynamics of tree growth, as it directly determines the capacity of trees to deliver ES over time (Elliot et al., 2019; Rötzer et al., 2021). Accurately modelling such dynamics requires explicit consideration of both spatial and temporal dimensions, which influence the way trees interact with their environment and, consequently, the way they function as service providers (Franklin et al., 2012; Moustakas et al., 2019).

When modelling urban trees, working at a sufficiently detailed spatial resolution is essential. Often urban trees are planted as isolated individuals or organized in linear formations (e.g., along streets or in small green patches) rather than in large continuous stands. In these settings, using coarse spatial grids becomes problematic because they fail to capture the heterogeneity of urban land cover, typically a fine-grained mosaic of impervious surfaces, vegetation patches, and built structures (Boehnke et al., 2022). While grid-based models offer advantages in terms of spatial exhaustiveness and simplicity, they assume internal homogeneity within each grid cell, which limits their accuracy when applied to spatially complex conditions (Seidl et al., 2012). To overcome this, an individual-based modelling approach may offer a more suitable alternative. It permits a more accurate representation of spatial heterogeneity, species-specific traits, localized disturbances, and site-specific management, making it a particularly effective approach in dense and fragmented urban environments (Lin et al., 2019).

An adequate representation of the temporal dimension is also fundamental when modelling urban trees and the ES they provide, since different ecological processes operate over distinct time scales (Almeida and Sands, 2016). For example, photosynthesis and evapotranspiration vary on an hourly basis in response to meteorological conditions, while biomass accumulation, leaf phenology, or carbon storage change more gradually over longer time scales. Selecting an appropriate time step for each process is therefore critical to maintain model realism, ensure sensitivity to environmental variability and thus ES provision (Duarte Rocha et al., 2022; Rau et al., 2020). Assuming that trees are static over time can lead to significant inaccuracies, especially when assessing their performance under non-stationary environmental conditions, such as those caused by climate change. In particular, low temporal resolution can obscure the effects of extreme events *e*.*g*., heatwaves and droughts, which often occur over short timescales but have disproportionately large impacts on tree physiology and service delivery (Cregg and Dix, 2001; Teskey et al., 2015). Similarly, long-term trends such as urban warming, increasing aridity, or altered seasonal dynamics cannot be captured without modelling growth as a dynamic and time-dependent process (Eickenscheidt et al., 2019; Esperon-Rodriguez et al., 2025). Therefore, to effectively support the planning of urban greening under future climate scenarios tree models should be structured to operate at high temporal resolution, to accurately simulate processes driven by radiation, temperature, and water availability.

Several models have been developed in recent years to estimate ES provided by urban trees and to simulate their growth under different conditions. These include widely used tools such as i-Tree (https://www.itreetools.org), which offers robust estimations of ES based on empirical inventories, as well as more spatially explicit frameworks like InVEST (https://naturalcapitalproject.stanford.edu/software/invest), which incorporate land use data and simplified vegetation modules. While these tools have greatly contributed to the operationalization of urban ES assessments, they are typically static or rely on average allometric relationships, which limits their responsiveness to dynamic factors such as extreme weather events, long-term climatic shifts or changes in management. Beyond these mainstream tools, a few process-based or hybrid models are emerging that seek to incorporate more environmental dynamism into the modelling of urban vegetation and their ES supply. Among the more recent modelling efforts, *CityTree* (Rötzer et al., 2019) represents an important step toward individual-based, process-oriented simulations of urban trees. It integrates photosynthesis, water balance, and stomatal regulation, allowing hourly resolution of transpiration and growth processes. Despite the many advanced features, the tree annual growth dynamic is *a priori* fixed, and thus the model less responsive to climate changes. A related model by Rötzer et al. (2021) focuses on interspecific variability in water use and drought response across different urban tree species, highlighting the role of physiological traits in ES delivery under stress. *UrbanTree* (Tams et al., 2023) builds on these foundations to simulate evapotranspiration and water stress dynamics in urban environments, considering the effects of shading and local microclimate on tree function. While it provides high temporal resolution, it does not include a dynamic growth component. In parallel, the model proposed by Babí Almenar et al. (2023) adopts a system dynamics approach to simulate the long-term evolution of urban trees and their ES supply, incorporating processes such as tree mortality, which is influenced by species drought tolerance, site conditions, and vegetation management. These models mark significant progress in the urban tree modelling, yet they remain limited in the capability of dealing with future climate scenarios. This highlights that there is still the need for integrated, modular, and climate-sensitive models tailored specifically to the complexity of urban forestry.

This paper presents a novel model, DynaTree, that adopts a mechanistic, dynamic, and individual-based approach to simulate the growth of urban trees and their provision of ES. The aim is addressed through three specific objectives:

i. to develop the conceptual and mathematical scheme of the model;
ii. to adapt the model to Milano, as a pilot case study, including calibration for three tree species and sensitivity analysis of key model parameters;
iii. to apply the model to assess the responses of the selected tree species across different planting ages and climate scenarios.

DynaTree is termed mechanistic because it represents the causal processes driving tree growth and ES supply, thereby avoiding exclusive reliance on empirical or statistical correlations. This approach enables the simulation of future conditions, such as those projected under climate change scenarios, without extrapolating beyond the domain of known data. The model is designed to offer a robust yet interpretable representation of key biophysical processes, to understand how tree growth and ES supply respond to environmental changes over time. It is also developed to be accessible, ensuring a simple interpretation and use by both expert modellers and practitioners in forestry, urban planning, and related fields.

## 2. Methods

### 2.1. Model scheme

As visually sketched in Figure 1, DynaTree is composed of a main core (Tree Biophysical Processes), which is informed by the inputs, and subsequently allows the calculations of ES in the dedicated module. This scheme reflects the logic of the ES cascade frameworks and the SEEA-EA accounting structure (United Nations et al., 2021). According to these frameworks, to monitor and measure the provisioning of ES, it is first necessary to model or gather data on the ecosystems’ biophysical structure, along with the underlying ecological processes and functions producing the services. Only after these steps have been accomplished can the supply of ES be estimated.

**Figure 1.**
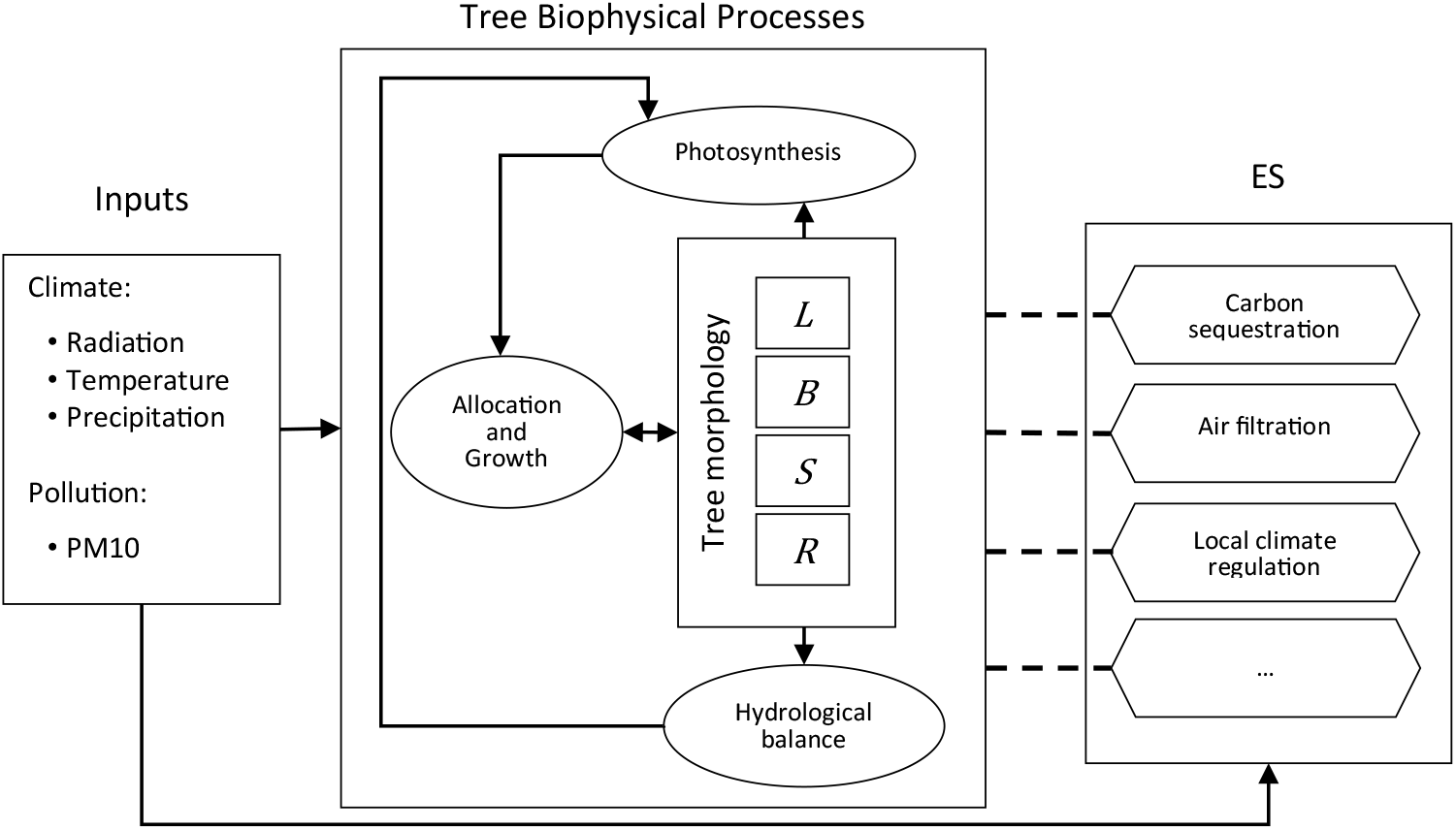
Schematic structure of DynaTree. Inputs (e.g. climate, pollution) feed into the Tree Biophysical Processes core, which simulates photosynthesis, carbon allocation and organs’ growth, hydrological balance, and tree morphology (by splitting the tree biomass into leaves, branches, stem, and roots). Outputs from this inform models for ecosystem services (e.g. carbon sequestration, air filtration, local climate regulation), which can be activated as needed.

The model is informed by inputs data of abiotic type, specifically meteorological and pollution series. For the calibration processes and for initializing the simulations also tree and soil data are needed (such as the tree Diameter at Breast Height (*DBH*), crown diameter (*CD*), tree age and species).

The Tree Biophysical Processes core describes the processes related to the growth of a single plant, and it is composed of four modules: photosynthesis, allocation of energy and biomass increase, hydrological balance and tree morphology. This last module does not represent a process in itself, rather it describes how the tree biomass distributed through time in each of the four model state variables: Leaves *L*(*t*), Branches *B*(*t*), Stem *S*(*t*) and Roots *R*(*t*). This splitting into multiple variables permits for example to model the biomass of leaves as linked to, yet separated from, the ligneous part or the tree, and thus to better describe the aboveground size of the tree, which is measured in terms of biomass of branches and stem.

DynaTree is complemented by an ES module, which take as inputs, together with the values of other parameters, also the values of the state variables computed through time *t* by the equations of the Biophysical Tree Processes. It comprehends different independently calculated ES, which can be added, activated and deactivated depending on the specific objectives of the analysis and the availability of data.

#### 2.1.1. Photosynthesis

The first module simulates tree photosynthesis, the primary driver of biomass accumulation and energy input into the system. Rather than describing photosynthesis at the biochemical level, DynaTree adopts a simplified, yet well established, empirically informed formulation that relates productivity to canopy surface and environmental constraints (Bahrami et al., 2022; Running et al., 2000; L. Zhang et al., 2015), through the equation:

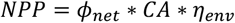

where the net primary production *NPP* [*kg*_*C*_/*day*] is estimated via the product of the specific net photosynthetic rate *ϕ*_*net*_ [*kg*_*C*_/*day*/*m*^2^], the crown area *CA* [*m*^2^], and a reduction factor *η*_*env*_ [−] capturing environmentally induced limitations on potential productivity of the tree. Ideally, *η*_*env*_ can depend on various environmental conditions (Pei et al., 2022) but, to minimize complexity, we will assume here dependence on water availability only, since that has been recognized to be the major limiting factor for tree growth in urban settings (Clark and Kjelgren, 1990; Dale and Frank, 2022). The quantitative method used to describe *η*_*env*_, as well as to assess the crown area *CA*, will be described respectively in Section 2.1.4 and Section 2.1.3. As for *ϕ*_*net*_, we calculated it as proportional, through various efficiency factors, to the incoming solar radiation *Rad* [*MJ*_*photons*_/*m*^2^]. Using the Beer-Lambert law, under the assumptions that both leaves and crown act as homogeneous media as in Landsberg and Waring (1997), we set:

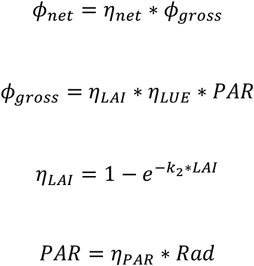

where *η*_*net*_ represents the net efficiency of primary production, whose value is later assumed equal to 0.45 [−] (Landsberg and Waring, 1997), *ϕ*_*gross*_ is the specific gross primary productivity [*kg*_*C*_/*day*/*m*^2^]. *PAR* represents the photosynthetic active radiation [*MJ*_*photons*_/*m*^2^], while *η*_*LAI*_ is the canopy radiation extinction term (Forrester, 2014; Natr and Lawlor, 2005), an adimensional factor [−] dependent on the leaf area index *LAI* in an increasing yet saturating way. The light extinction coefficient regulating saturation *k*_2_ is an empirical parameter whose value is usually set in the range 0.4-0.7 (Natr and Lawlor, 2005; Smith, 1993) and calibrated in this study making use of data from Milano (see Section 2.2). *η*_*LUE*_ is the light use efficiency [*kg*_*C*_/*MJ*_*photons*_], and its value is set to 0.0036, as derived from the value of 0.063 [*mol*_*C*_/*mol*_*photons*_] found in the database for tree traits TryDB (Kattge et al., 2020) (Table S1, Supplementary Material). Lastly, *η*_*PAR*_ is the part of radiation usable by the tree, equal to 0.5 [−] (Landsberg and Waring, 1997).

#### 2.1.2. Allocation and Growth

The second module governs biomass allocation and loss, providing a simplified representation of how *NPP* is dynamically distributed across tree compartments. The structure of this module is based on a set of differential equations describing biomass accumulation and depletion over time in each relevant tree compartment *i*, elaborated over those presented in Bevacqua et al. (2021):

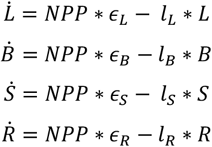

The dimensionless parameter *𝜖*_*i*_ represents the allocation of *NPP* to the compartment *i* (as detailed below) while the parameter *l*_*i*_ [1/*day*] is the compartment dependent decay rate regulating the biomass loss that accounts for the reduction of biomass in the tree compartment *i*. The energy entering through photosynthetic carbon uptake in the system via *NPP* is dynamically allocated to each compartment (*L, B, S* and *R*) and is outgoing the system proportional to the stock of biomass in the compartment.

The allocation coefficients *𝜖*_*i*_ are dimensionless and defined as follow:

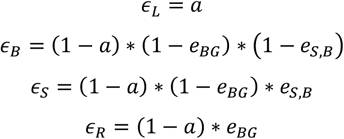

The parameter *a* regulates seasonal energy allocation to leaves. Since no clear references exist, we adopted a simplified formulation to approximate the natural dynamics of broadleaved deciduous trees. A low baseline (*a* = 0.01) is assumed most of the year, increasing in spring (*a* = 0.5) to mimic budburst and then declining, capturing the essential phenological pattern. The timing is determined by a budburst function based on temperature accumulation (Malyshev et al., 2024; Murray et al., 1989; Prentice et al., 1992)(further explained in Supplementary Material, Appendix A). The belowground allocation coefficient *𝜖*_*BG*_ can vary between 0.23 and 0.8 depending on tree stress level, as reported by Landsberg and Waring (1997). For Milan, we estimated an average value of 0.3 under typical annual conditions, which was adopted as the fixed coefficient in the model. The parameter governing the allocation split between stem and branches *e*_*S,B*_, cannot be directly desumed from equivalents available in the literature. However, reported stem-to-branch biomass ratios typically range from 2.5 to 3.5 (He et al., 2023; Martin et al., 1998). The parameter was thus not directly fixed but empirically calibrated, constraining it between 0 and 1 (as it represents the fraction of biomass allocated to each compartment), so that the resulting model outputs respected stem-to-branch biomass ratios range during the calibration phase.

The loss coefficient *l* varies by compartment. For leaves, *l*_*L*_ includes a baseline of 0.1% per day (Meier et al., 2006), representing a minimal continuous loss. For temperate deciduous species, *l*_*L*_ is increased in autumn to simulate leaf fall, reducing leaf biomass to near-zero (full details in Supplementary Material, Appendix A). For stem and branches, no robust values of loss coefficient were found in the literature, thus, the loss coefficients *l*_*S*_ and *l*_*B*_ were calibrated (as visibile in Section 2.2). Instead, for roots, the turnover coefficient *l*_*R*_ was set to five times the value of *l*_*S*_, based on internal model consistency and intermediate validation tests. Although this represents a simplification, it is justified because root biomass does not directly affect tree growth in the current model.

#### 2.1.3. Tree morphology

This module consists of equations that translate the internal biomass dynamics into morphological attributes. These attributes are indeed essential to link physiological processes with ES supply, as they mediate, for example, light interception or pollutant filtering capacity. The identified relationships are species-specific and are derived either from geometric assumptions based on tree morphology or from allometric functions.

The calculation of the crown projected area, abbreviated as *CPA* and measured in [*m*^2^], is based on the relationship between branch biomass (*B*) and crown extension. Following empirical studies on tree architecture (Drew and Flewelling, 1977; Kuuluvainen, 1991), we can use the classical power-law formulation with a 2/3 exponent:

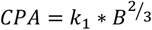

where *k*_1_ is a species-specific parameter calibrated so as to best fit available data.

To obtain the crown area (abbreviated as *CA*), we can multiply *CPA* by a form factor (*ff*) that accounts for the species-specific crown shape:

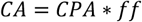

Crown shapes used in this study were classified as expanded, round, oval, pyramidal or fastigiate following an Italian urban tree catalog (https://ambiente.regione.emilia-romagna.it/it/radiciperilfuturoer/alberi-per-la-citta), assigning the following form factor values: 1 for round and expanded crowns, 1.25 for oval and pyramidal shaped crowns, and 1.5 for fastigiate crowns. *CPA* is typically assumed to be circular, thus the crown diameter *CD* is estimated as the diameter of a circle with area equal to *CPA*.

Leaf area *LA* [*m*^2^] and Leaf Area Index *LAI* [−] are respectively calculated as:

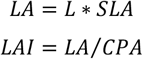

in which *SLA* refers to the specific leaf area, measured in *m*^2^/*kg*_*dry matter*_, sourced from TryDB database (Kattge et al., 2020) with the values available in Table S2 (Supplementary Material). *SLA* values are then converted from dry matter to carbon-based units using a fixed ratio of 2 *kg*_*dry matter*_./*kg*_*C*_ (De Vries et al., 2006; Keith et al., 2009). *LAI* maximum value is then set to 5 (Kimm and Ryu, 2015; Moser-Reischl et al., 2025), to ensure reasonable values and prevent overestimation of light interception.

Above-ground biomass (*AGB*) is considered to be equal to *S* + *B*, to consider only the woody compartments. Diameter at Breast Height (*DBH*) is estimated from the following literature derived allometric equations (Annighöfer et al., 2012; Fortier et al., 2017; McPhearson et al., 2013):

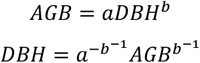

where parameters *a* and *b* are derived from literature and their values are reported in Table S3 of Supplementary Material. Even though the overarching conceptual framework of the model aims to minimize the reliance on empirical equations, allometric functions are employed here as a bridge to translate biomass into measurable structural dimensions of the tree.

#### 2.1.4. Hydrological balance

This module quantifies the water availability in the soil so as to set one of the most important abiotic conditions that determine whether the tree is growing under optimal conditions, experiencing heavy stress due to water scarcity.

The soil water dynamic is here described through a simplified hydrological balance equation:

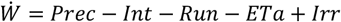

where *W* is the water level at time t [*mm*], *Prec* is the precipitation [*mm*/*day*] directly derived from data, *Int* is the interception [*mm*/*day*], *Run* is the Runoff [*mm*/*day*], *ETa* is the actual evapotranspiration [*mm*/*day*] and *Irr* is the irrigation [*mm*/*day*]. *Irr* can be manually changed to represent the management option adopted for the tree. *Int* is calculated as fixed percentage *p*_1_ of *Prec*, and in the same way *Run* is a fixed percentages *p*_2_ of of the precipitation remaining after interception. For simplicity, both *p*_1_ and *p*_2_ are set to 10%, providing a consistent approximation of water availability for the tree. Although these fixed values underestimate those reported in the literature (Cai et al., 2024; Xiao and McPherson, 2011; Zabret and Šraj, 2019), they provide a consistent and simple approximation of water availability for the tree. When the water level *W* exceeds the total available water *TAW* (further explained in Supplementary material, Appendix A), its level is set back to *TAW*. In general, this approach was taken to maintain model simplicity, as the goal is not to simulate detailed soil hydrology but to approximate the water available for the tree.

*ETa* is calculated as described in the Crop evapotranspiration Guidelines of FAO (Allen et al., 1998) as:

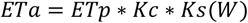

in which *ETp* is the potential evapotranspiration [*mm*/*day*], *Kc* is the crop coefficient [−] and *Ks* is the water stress coefficient [−] that is dependent on the water in soil *W*. More precisely, *ETp* is calculated according to the Hargreaves-Samani method (Hargreaves and Samani, 1985); *Kc* is dynamically described in the same way as *η*_*LAI*_, as described also in Chapter 9 of FAO Guidelines (Allen et al., 1998).

*Ks* is instead the reduction of the evapotranspiration dependent on water scarcity (Allen et al., 1998) and it’s calculated as:

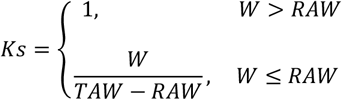

in which *TAW* is in [*mm*], and *RAW* readily available water [*mm*], which is the portion of *TAW* in which the water is free to move and so to be uptake from the plant (Supplementary Material, Appendix A). The stress coefficient *Ks* is then directly used to compute the environmental modifier of photosynthesis *η*_*env*_. This means that the photosynthetic rate decreases linearly from optimal (*η*_*env*_ = *Ks* = 1) to null (*η*_*env*_ = *Ks* = 0) as water availability drops below critical thresholds.

#### 2.1.5. Ecosystem Services

The ES module, which can in principle account for various other services, is composed here of the following three ES: carbon sequestration, air filtration and local climate regulation. These were selected due to their primarily relevance in enhancing urban livability, and in contrasting and mitigating climate change (Babí Almenar et al., 2021; Castellar et al., 2021; Li et al., 2025). Each ES sub-module is directly fed by variables computed through time to account for biophysical processes of the tree, with outputs aggregated over time whenever needed.

Carbon sequestration is estimated as a direct function of *NPP*, expressed in *kg*_*C*_/*day*. The total carbon sequestered from the atmosphere is obtained by simply integrating *NPP* over the simulation period of interest.

For air filtration, the model just consider dry deposition of PM_10_, which can be calculated according to the resistance analogy method for PM_10_ (Manes et al., 2016; Nowak, 1994). The formula used here is derived from formulation of Babí Almenar et al. (2023):

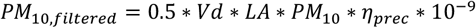

where *PM*_10,*filtered*_ is the flux of PM10 deposited on leaves in 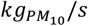, *Vd* is deposition velocity [*m*/*s*], *PM*_10_ is the concentration of particulate 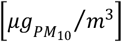 and *η*_*prec*_ is a boolean factor to skip wet deposition during rainy days (e.g. those when precipitation is above 1 *mm*). As for *Vd* is parameterized here as in Yang et al. (2005):

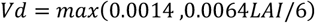

Local Climate Regulation is defined as proportional to the actual evapotranspiration of trees, through factors that converts the amount of evapotranspirated water into kilowatt-hours (kWh) of energy-equivalent cooling (Babí Almenar et al., 2023; Moss et al., 2019). Following the amending EU Regulation on environmental economic accounts (European Union, 2024), to account for days in which local climate regulation is relevant, the maximum temperature must be above 25 degrees. To convert evapotranspiration values into energy savings, defined in kWh, the following formula is used:

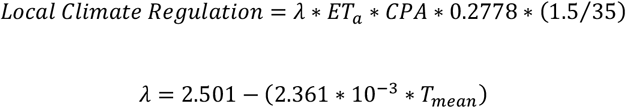

where *λ* represents the latent heat of evaporation [MJ/kg], which varies with the mean daily temperature (*T*_*mean*_).. The factor 0.2778 is the conversion factor from megajoules (MJ) to kilowatt-hours (kWh), while the ratio 1.5/35 reflects the coefficient of performance of the reference air conditioning system, indicating that 1.5 kWh of electricity is required to provide 35 kWh of cooling. This air conditioning performance reference is consistent with the one used in Moss et al. (2019), and should be progressively aligned with the efficiency of man-made cooling systems.

### 2.2. Calibration and sensitivity analysis

The parameters subject to calibration were those lacking reliable literature values or considered to be largely empirical:

- *k*_1_: the correlation factor between *B* and *CPA*;
- *k*_2_: the light extinction coefficient;
- *l*_*S*_: the stem biomass loss;
- *l*_*B*_: the branch biomass loss;
- *e*_*S,B*_: the mass allocation coefficient between branches and stem.

The calibration process was performed using the data obtained from the Open Data portal of the Municipality of Milan (Comune di Milano, 2023)(further explained in Supplementary Material, Appendix B), precisely on *DBH* and *CD* values.

To smooth out gaps in the annual age distribution of trees, we group tree data for both *DBH* and *CD* into 5-year age classes, starting from 3 years old trees [3-7, 8-12, …]. For each class, the third quartile (Q3) was taken as a rappresentative value for a healthy, well-performing tree. Calibration of the unknown parameters was then conducted using the genetic algorithm NSGA-II (Deb et al., 2002), to simultaneously minimize two objective functions, namely the mean square error between the model predictions and the third quartiles for both *DBH* and *CD*. In order to assess whether the properly calibrated model had an acceptable predictive capability, we halved the original dataset into two subset, a calibration subset (with 50% of the data) and a validation subset (with the remaining 50%).

Since the calibration is multi-objective, its outcome will not be a vector of optimal parameter values. Instead, it yields a set of solution vectors constituting in the two-dimensional objective space, forming the so-called Pareto frontier. To evaluate model robustness, we considered all Pareto-optimal solutions during the calibration analysis, which allowed us to assess the range of plausible parameter combinations. For the pilot application across the three climate scenarios, however, we selected a single solution for each species corresponding to the utopia point (i.e., the point on the Pareto frontier closest to the origin) representing the best compromise between the two calibration objectives. This solution was then used in all subsequent simulations.

To evaluate the sensitivity of DynaTree to parameter variation, we performed a one-at-a-time (OAT) sensitivity analysis (Saltelli, 2008) on the five calibrated parameters: *k*_1_, *k*_2_, *l*_*S*_, *l*_*B*_ and *e*_*S,B*_. All parameters were individually perturbed by ±20% around their calibrated values. During the sensitivity analysis all the other parameters were held constant at their resulting optimal values. The ultimate goal of these perturbations was to assess how variations and/or possible uncertainties in each parameter may influence the model’s predictions of tree growth and ES supply.

For each parameter perturbation, the resulting tree growth trajectories were plotted and compared against the baseline simulation, so as to visually compare which parameter variations most strongly influence model outcomes and whether the system behaves robustly under plausible uncertainty.

### 2.3. Case study and climate change scenarios

The city of Milan was selected as a case study. From the tree data of Milan dataset (Supplementary material, Appendix B), we selected three of the most common tree species in the city for a pilot application of the model: *Platanus × acerifolia, Populus nigra* and *Robinia pseudoacacia*. These species also differ in their biogeographic status, i.e., whether their presence is considered native or alien. *Populus nigra* is autochthonous in Milan, *Platanus × acerifolia* is a man-made hybrid naturalized in the city, and *Robinia pseudoacacia* is an invasive species.

To assess DynaTree’s performance for the next 25 years (i.e. 2025-2049) under different climatic conditions, we selected three climate scenarios that represent contrasting future trajectories: a historical baseline (i.e. the BAU scenario), the very stringent pathway to mitigate emission (RCP2.6 mitigation scenario), and the worst-case pathway (RCP8.5 high-emission scenario). These scenarios were chosen to explore a wide range of possible future climates, from low-impact to extreme warming conditions, and to test the model’s sensitivity and suitability for projecting urban tree growth and ES supply beyond historical baselines. The BAU scenario serves both as a reference and as a calibration baseline, while RCP2.6 and RCP8.5 provide distinct, somehow opposite alternatives for evaluating the potential impacts of climate change on urban tree performance over time under two very different development hypotheses. The meteorological data were sourced from the CORDEX regional climate model ensemble (Copernicus Climate Change Service, 2019)(for the specification see Supplementary Material, Appendix C).

As for air pollution, PM_10_ concentration data were retrieved from an ARPA Lombardia monitoring station in Milan (ARPA Lombardia, 2024) which provided 10 years of daily measurements (2011–2020). Based on these records, we generated a synthetic daily time series with equivalent statistical properties, extended to match the full simulation period required by the model (Supplementary Material, appendix C). Also, in future scenarios we use do not consider changes in the PM_10_ concentration.

Using the calibrated parameter sets, we simulated the growth of each species starting from initial ages of 5, 10, and 20 years for the 25-year period under all three climate scenarios. These simulations allow us to explore how tree growth and ecosystem service provision evolve over time under different environmental conditions, reflecting both species-specific traits and the effects of climate forcing. By including multiple starting ages, the model captures differences in growth trajectories and ES delivery across life stages, providing insights for urban forest planning, maintenance strategies, and long-term management of ecosystem services in Milan.

## 3. Results

### 3.1. Calibration and sensitivity analysis

The calibration procedure identified a two-dimensional set of Pareto-optimal solutions that minimize the combined error in predicting *DBH* and *CD* of the tree database for Milan. Figure 2 shows the resulting optimal trade-offs (top row) for *Platanus × acerifolia, Populus nigra* and *Robinia pseudoacacia* along with the simulated growth trajectories of *DBH* (second row) and *CD* (third row) associated with the selected parameter set. Each point of the Pareto frontier corresponds to a unique combination of parameters, revealing how the improvement of one objective results in comprosing performance in the other.

**Figure 2.**
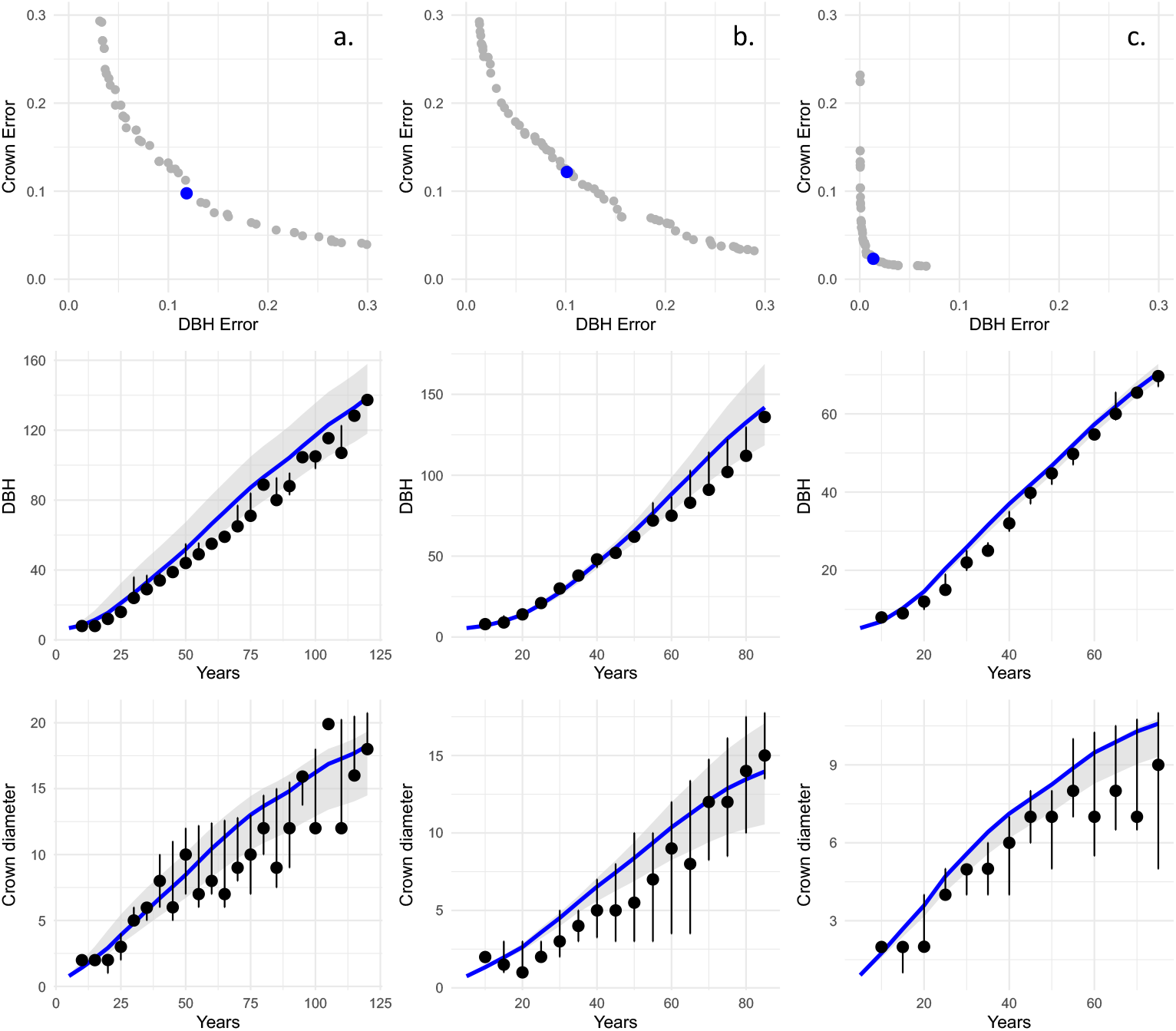
Calibration of our model on tree data of the Municipality of Milan, with reference to the three species (a. Platanus x acerifolia, b. Populus nigra, c. Robinia pseudoacacia). Top row: Pareto frontiers showing the trade-off between DBH and crown diameter error for each species; errors values refers to average squared deviations in standardized units; the blue dot indicates the selected optimal solution to perform simulations in the other panels. Middle and bottom rows: Simulated DBH and CD growth curves (blue lines) for the corresponding selected parameter set, contrasted to the empirical values in the dataset grouped by age class (black dots, median ± IQR). Shaded areas represent the range of model trajectories within the pareto set parameter variation.

The Pareto frontiers in the top row of Figure 2 highlight distinct calibration patterns across species. *Platanus × acerifolia* and *Populus nigra* display broadly similar frontier shapes, with comparable ranges of *DBH* and *CD* errors. These errors are expressed as average squared deviations in standardized units, i.e., errors are normalized by the variance of observed values. In contrast, *Robinia pseudoacacia* shows a much more compact frontier, with errors of nearly an order of magnitude lower and far less dispersed, particularly for the DBH error. This suggests both a better overall model fit and a reduced variability in the data used for calibration. The tighter clustering of points could be attributed to a more homogeneous growth behavior of *Robinia* individuals.

In terms of fit, the second and third rows in Figure 2 reveal that for *Platanus* there is a general good agreement between the model and the empirical data. For *DBH*, there is a slight tendency to overestimate values in later life stages of the plant, while for *CD*, the fit is a bit less precise, partly because of the high variability and irregular distribution of the empirical data, which may reflect limitations in measurement or inconsistencies in data quality. *Populus* displays a very good fit for *DBH* throughout the tree lifespan, with model predictions closely tracking observed values. For *CD* data the model captures the general trend with the only tendency to underestimate crown size for older individuals. This may reflect higher uncertainty in the observed values, likely due to the visual estimation method used by different trained operators in early age classes, which is inherently more subjective than DBH measurements. *Robinia* stands out for the overall quality of fit. Both *DBH* and *CD* are well captured by DynaTree, with very limited deviation from the observed data. This strong agreement reinforces the idea that Robinia’s growth pattern is more consistent across individuals and better aligned with the model’s structural assumptions.

Overall, the calibration results confirm that DynaTree is capable of reproducing realistic growth trajectories across species, even in the presence of noisy data. The goodness of fit varies between species, both because of species-specific characteristics and of data quality, but our results provide a solid foundation for applying the model under present conditions and future climate scenarios.

Table 1 reports the values of the calibrated parameters corresponding to the utopia points. The parameter *k*_1_, which regulates the relationship between branch biomass and crown projected area, shows a relatively wide range of variability across the selected species, reflecting interspecific differences in crown expansion, further illustrated in Figure S1 (Supplementary Material). In contrast, the parameter *k*_2_, which defines the relationship between leaf biomass and light interception, displays small variations across species, especially between *Populus* and *Robinia*. The parameter regulating the loss of biomass (and so the turnover) of branches *l*_*B*_ is consistently about one order of magnitude higher than the loss rate of stem *l*_*S*_ for all species, aligning with the idea of faster biomass turnover in peripheral compartments (H. Zhang et al., 2015). Lastly, the allocation parameter *e*_*S,B*_, which determines the partitioning of biomass between stem and branch compartments, shows a very narrow range of variation, suggesting a relatively stable allocation strategy across the three species studied.

**Table 1:** Calibrated parameter values for our model corresponding to the utopia point for each species. The parameters include morphological coefficients (k_1_, k_2_), turnover rates (l_S_, l_B_), and the allocation coefficient (e_S_) governing biomass partitioning between stem and branches.

| Tree species | $k_1$ | $k_2$ | $l_S$ | $l_B$ | $e_{S,B}$ |
| --- | --- | --- | --- | --- | --- |
| *Platanus x acerifolia* | 1.712 | 0.452 | 1.04E-05 | 1.01E-4 | 0.632 |
| *Populus nigra* | 1.528 | 0.342 | 1.94E-05 | 1.61E-4 | 0.609 |
| *Robinia pseudoacacia* | 2.138 | 0.356 | 3.80E-05 | 3.50E-4 | 0.557 |

The results of the sensitivity analysis, in which the five calibrated parameters were varied within a ±20% range around their calibrated value, confirm the overall robustness of the model across species and parameters. The species-specific impacts of parameter variation in the predicted trajectories of *DBH* and *CD* are illustrated in Figure 3.

**Figure 3.**
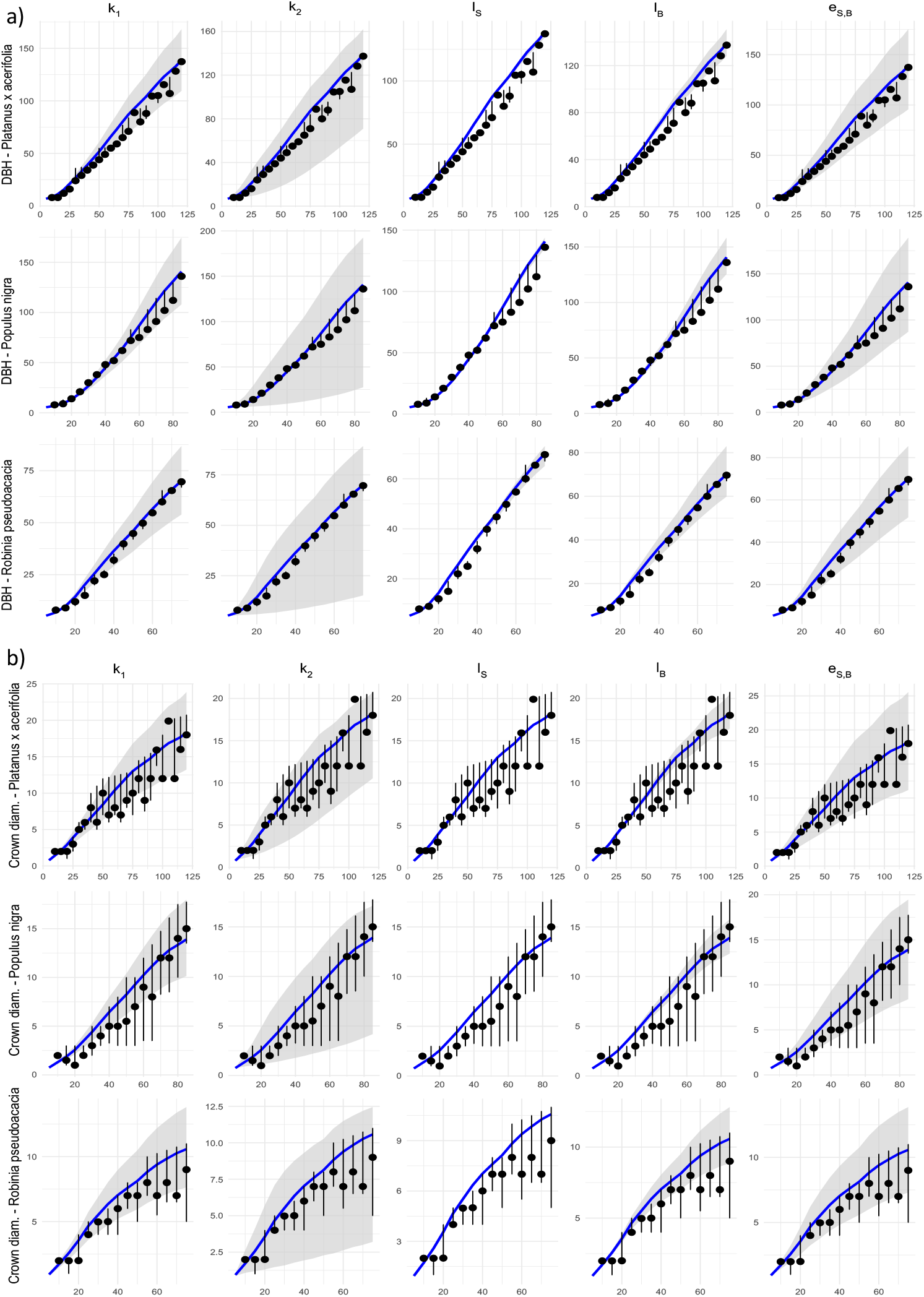
Sensitivity analysis for the five calibrated parameters: k_1_, k_2_, l_S_, l_B_ and e_S,B_. For each parameter, DBH (top, group of panels a) and CD (bottom, group of panels b) projections are shown under ±20% variation (gray bands) around the baseline simulation (blue curve). Black dots and error bars indicate the empirical quartile values per grouped age classes used for model calibration.

Despite the variation in parameter values, trees continue to grow following biologically plausible trajectories, and no parameter variation produces unrealistic behaviors or model breakdowns. Nonetheless, the impact of individual parameters differs substantially by species and by target variable. Among the tested parameters, *k*_*2*_ and *e*_*S,B*_ seem to exert the strongest influence on model outputs, particularly for *Robinia* and *Populus*, with changes that peak at a 70% of difference. In contrast, *k*_*1*_ induces more moderate, but still significant deviations, especially for crown projections (with differences that peak at around 30% of the projected values).

Biomass loss parameters behave differently. The effect of perturbations to *l*_*S*_ appear consistently low across species, with deviations that are almost negligible. While this analysis suggests that *l*_*S*_ could be fixed a priori to restrict the set of parameters to be calibrated, we opted to retain it as a calibrated parameter due to its high interspecific variability. *l*_*B*_ has a larger influence on projections, but remains consistent with the original projections, with deviations that do not exceed 15% (the parameter perturbation propagates less than linearly to results).

### 3.2. Model functioning and growth under climate scenarios

To understand DynaTree’s responses to future climates, we first examine its behavior under baseline conditions. Figure 4 illustrates the dynamic allocation of biomass, a core aspect of DynaTree’s internal functioning. Each species shows a progressive accumulation of structural biomass in stems and branches, while leaf biomass exhibits a distinct seasonal pattern. The model successfully captures this cyclical behavior, which aligns with the known phenology of temperate species.

**Figure 4.**
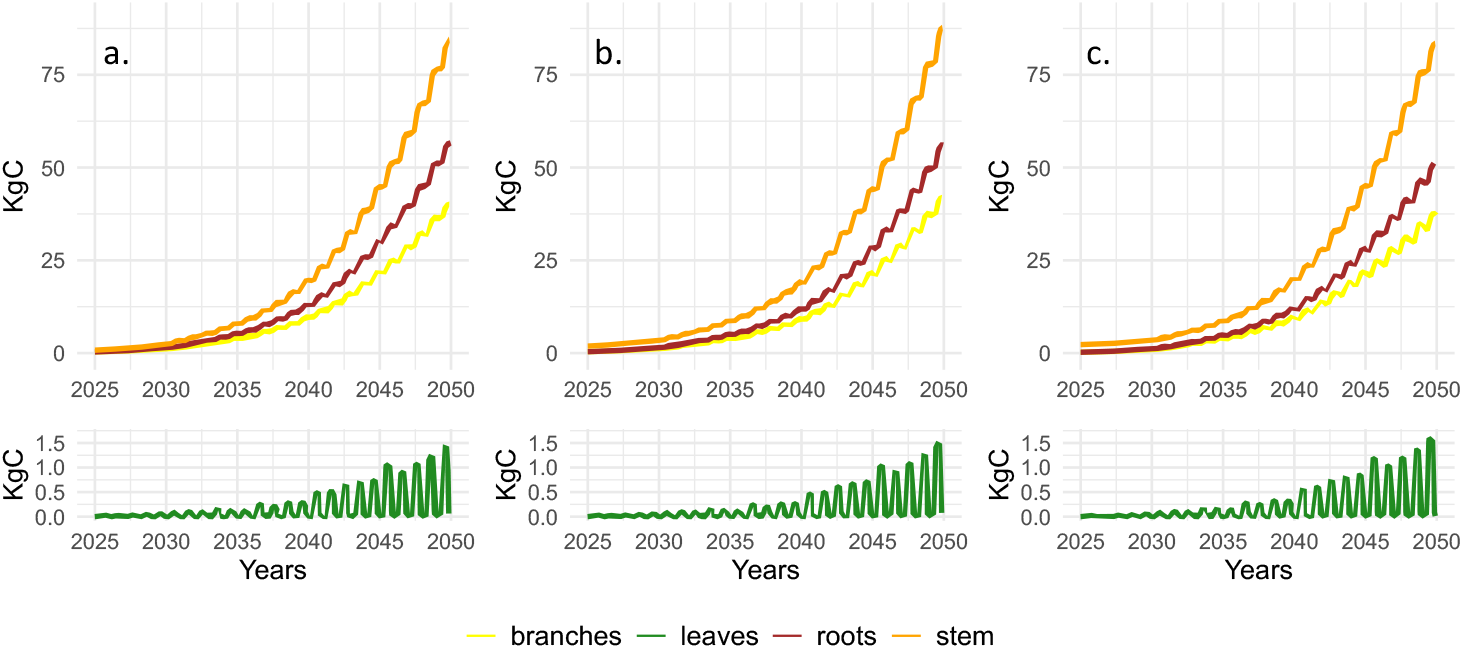
Simulated biomass allocation between plant compartments for the three focal species (a. Platanus x acerifolia, b. Populus nigra and c. Robinia pseudoacacia) under historical climate regimes for 5-years-old trees planted in 2025. Each panel in the top row shows the biomass dynamics of stems (orange), branches (yellow), and roots (brown), while in the bottom row is leaf biomass (green). Seasonal fluctuations in the leaf compartment highlight intra-annual growth and senescence dynamics for the deciduous species.

Once the dynamics of biomass allocation between tree compartments has been clarified under current climate conditions, it is useful to explore future scenarios possibly assessing the impact of climate (baseline historical conditions, RCP 2.6 and RCP 8.5). Figure 5 shows the total biomass accumulated in all compartments, expressed in carbon units (kgC), over the 2025–2050 period for the three species and starting ages. This allows us to compare how different climate forcings (color coded in Fig. 5) affect trees of different ages (linestyle coded) so as to shape tree growth trajectories under various environmental conditions, in particular temperature and precipitation.

**Figure 5.**
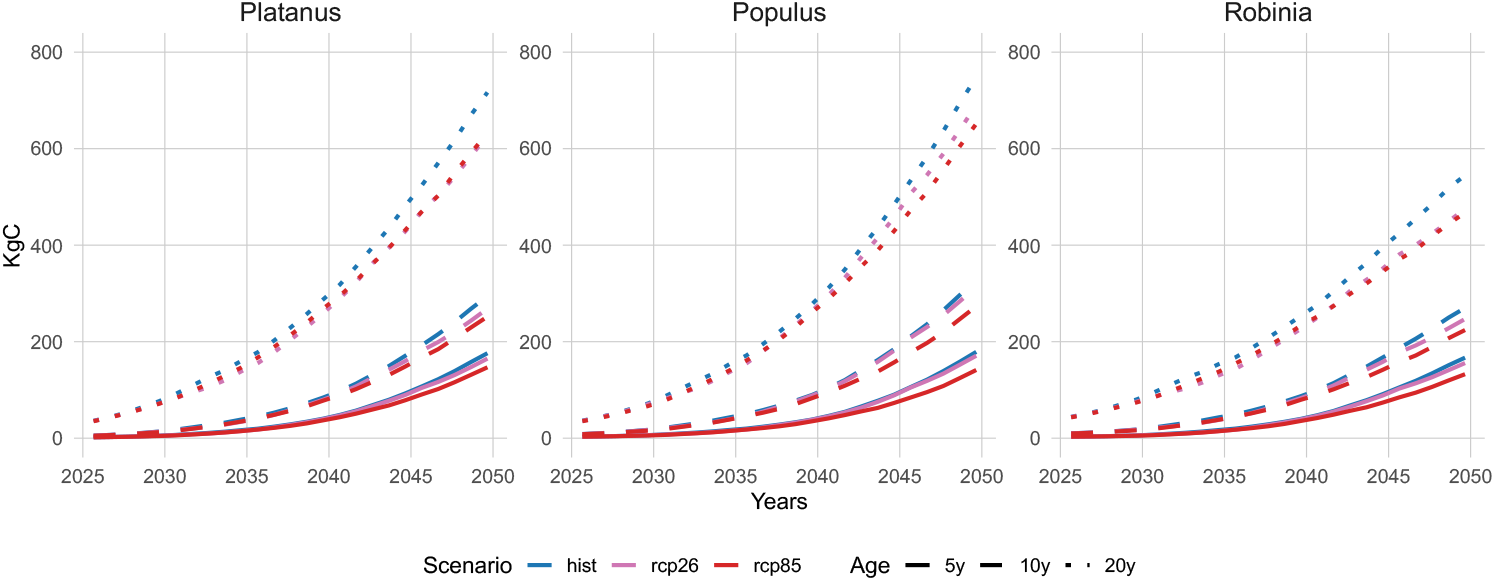
Projected total biomass accumulation (kgC) for the three species (in the different panels) under three climate scenarios (Historical in blue, RCP2.6 in pink, RCP8.5 in red) and three initial ages (5 years plain lines, 10 years dashed lines, 20 years dotted lines).

Across all species and scenarios, clear patterns emerge. Biomass accumulation decreases with increasing climate forcing: for all species, the RCP8.5 scenario leads to noticeably reduced growth compared to the historical scenario, while RCP2.6 shows intermediate values. Despite this general trend, in terms of species-specific impacts of climate change on urban trees, according to our model applied to the case of Milan, *Populus* appears slightly more resilient to climate forcing, with smaller differences among scenarios than the other two species. Intriguingly, the species-specific response to RCP2.6 seems to be different depending on age: for younger trees, the growth pattern under RCP2.6 is nearly identical to that occurring under unchanged climate, but as tree age increases, RCP2.6 curves tend to be more similar to the RCP8.5, especially in the case of *Platanus*. This suggests that the effects of moderate climate change may become more pronounced over longer lifespans or for larger individuals.

Our analysis shows that DynaTree can help to discuss interspecific and age-dependent responses to climate, indicating that species traits and initial conditions both modulate growth under future scenarios.

### 3.3. Ecosystem services scenarios

The performance of the three species for carbon sequestration (Figure 6), air filtration (Figure 7), and local climate regulation (Figure 8) across climate change and management scenarios (age of planted trees) is summarized in the following lines.

**Figure 6.**
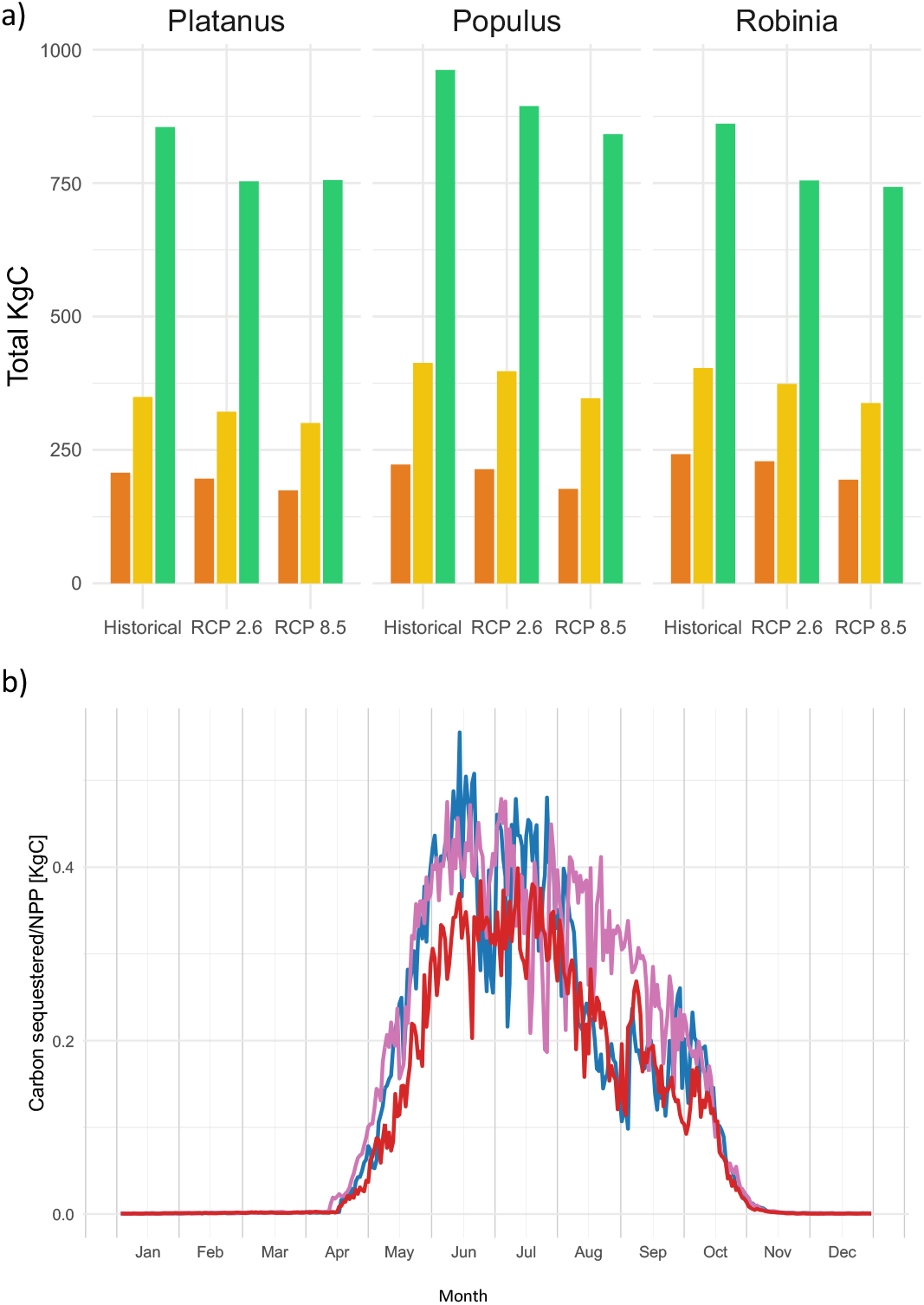
Model computation of the carbon sequestration ecosystem service. a) Total carbon sequestered over the simulation period 2025-2050 for each of the three species, according to the climate scenarios, and for three different cohorts of trees (aged respectively 5 years in orange, 10 years in yellow and 20 years in green). b) Daily carbon sequestration (equal to NPP) during one year (we used 2044) for a 10-years old Populus tree; colors correspond to the different climate scenarios as in Figure 5.

**Figure 7.**
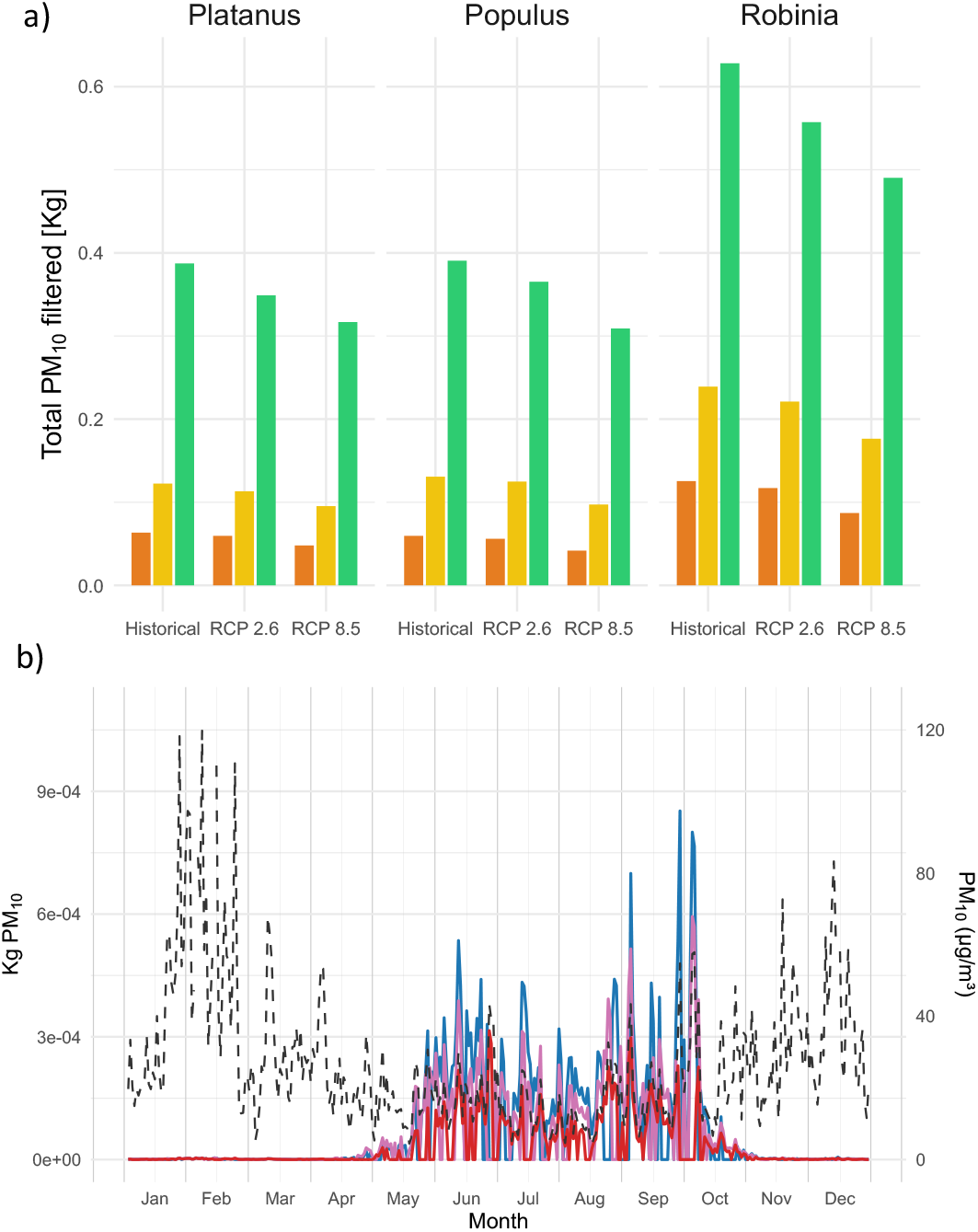
PM_10_ filtration. a) Total PM_10_ removed during the 2025-2050 period by trees of different species and ages (aged 5, 10, 20 years in 2025, respectively in orange, yellow and green) under different climate scenarios. b) Daily filtration for a 10-year-old Robinia in 2044, under the different climate scenarios (Historical, RCP 2.6 and RCP 8.5, respectively in blue, pink and red), with PM_10_ concentration shown in dashed black line (the level are visible on the right axis).

**Figure 8.**
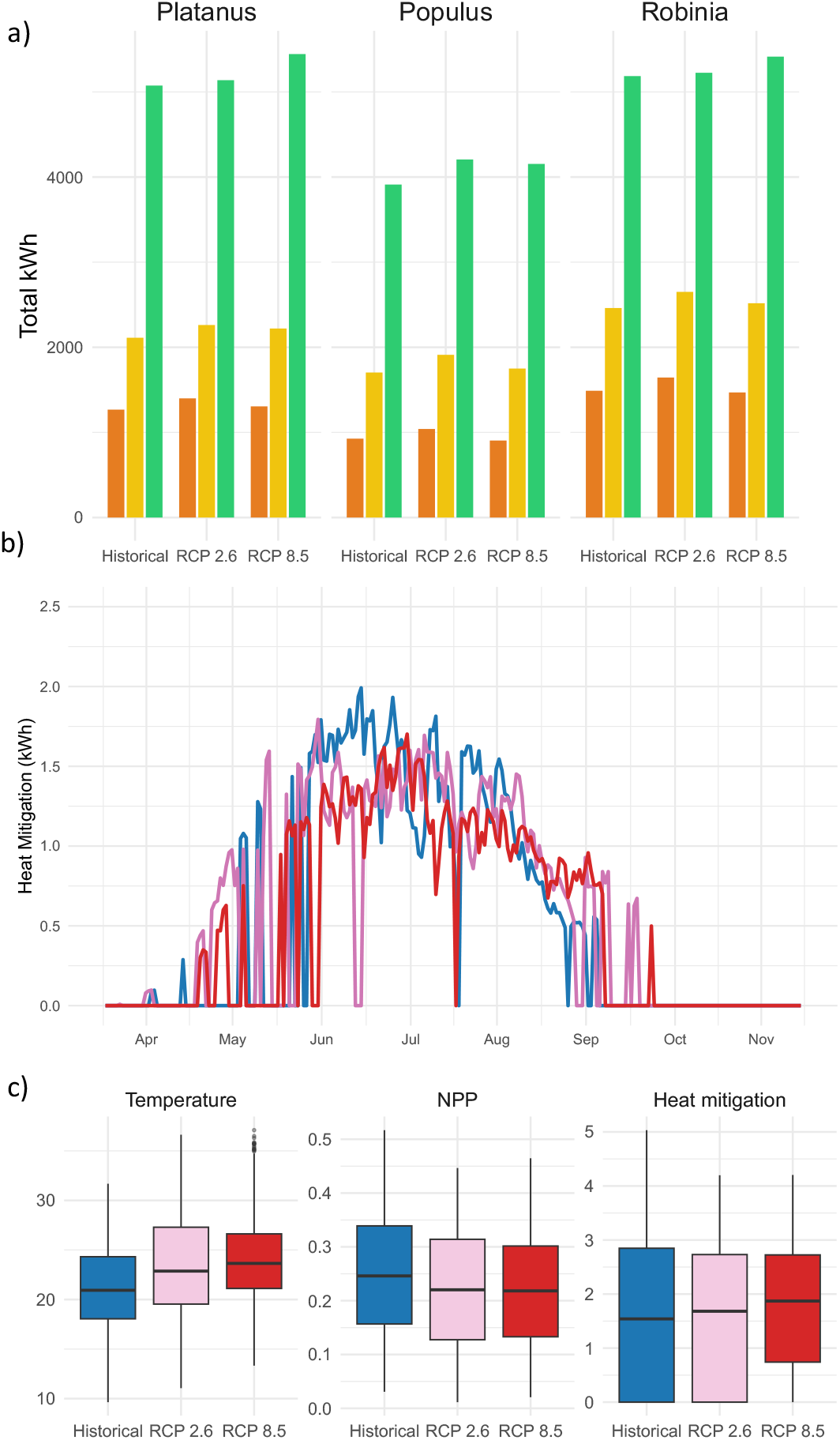
Heat mitigation. a) Total energy-equivalent cooling (in kWh) provided by trees under different scenarios and initial ages (orange for 5 years, yellow for 10 years, green for 20 years). b) Daily heat mitigation in an exemplificative future year (here is 2041) for a 10-year-old Platanus under the three climate scenarios (Historical in blue, RCP 2.6 in pink, RCP 8.5 in red). c) Boxplots of temperature, NPP, and total heat mitigation from the 5-years period 2045–2049 for a 10-year-old Platanus under the three climate scenarios.

In terms of carbon sequestration, as shown in Figure 6a, planted at the age of 20 years trees can deliver a disproportionately higher carbon uptake over the simulation period compared to younger trees. This emphasizes the nonlinear accumulation of carbon with tree maturity. Figure 6a also shows a clear reduction in carbon sequestration under increasing climate forcing, particularly under the high-emission scenario RCP8.5, and such effect becomes more evident in more mature trees. Figure 6b complements this result by showing as an example the daily carbon net uptake from the atmosphere (which is in fact NPP) during a year (here is 2044) for a Populus tree aged 10 at the start of the simulation. While all scenarios follow a similar seasonal pattern, the RCP8.5 curve consistently lies below the others during the peak photosynthetic period, indicating reduced productivity under more stressful climatic conditions.

Regarding air filtration, Figure 7a clearly shows that *Robinia pseudoacacia* consistently outperforms the other species in terms of PM_10_ filtered, likely due to a higher LAI that enhances pollutant capture. As it occurs with carbon sequestration, a reduction in PM_10_ filtered is observed under more intense climate scenarios, particularly visible in mature trees. Figure 7b depicts daily PM_10_ filtration during 2044 for a 10-year-old Robinia tree. Even in this case, as it was the case for carbon assimilation, it is possible to appreciate a reduction of PM_10_ deposition as the climate forcing increases. This ES closely follows the pattern of PM_10_ concentration (dashed black line), with filtration peaking during periods of lower pollution. This occurs because of the evidence, captured by the model, that dry deposition of PM_10_ is proportional to leaf area and pollutant concentration and highlights that filtration occurs when the photosynthesis is more active, in spring and summer, that is when pollution is also lower in Milan.

In terms of heat mitigation, Figure 8a shows the cumulative values for the different trees and scenarios for the period 2025-2050. Unlike the other two ES, local climate regulation does not decline under stronger climate scenarios; values are stable or even slightly increased under RCP2.6 and RCP8.5 compared to historical scenarios. To deepen our understanding on this unexpected result, in Figure 8b we examine the service offered by 10-year-old Platanus tree in the year 2041, a future year when local climate regulation appears from our model to be very similar across scenarios (183 kWh for historical, 196 for RCP 2.6 and 184 kWh for RCP 8.5). As expected, the graph does not highlight any significant difference among the scenarios. Boxplots in Figure 8c (years 2045–2049) reveal the trade-offs that cause homogeneity across scenarios: while NPP decreases slightly under more intense climate scenarios, lowering tree biomass, temperature increases enhancing water loss by leaves. Since evapotranspiration, and hence cooling, is driven by both, the effect of declining tree vitality is offset by higher evaporative demand, resulting in overall similar local climate regulation among scenarios.

## 4. Discussion

### 4.1. Conceptual and Mathematical Structure of the Model

Unlike many existing tools which rely on static inventories, empirical averages, or cell-based approximations, DynaTree explicitly represents causal processes operating at the tree level and over time. This allows for a more realistic simulation of how individual trees will respond to accelerated changes in environmental drivers, such as temperature and water availability, as expected under future climate scenarios.

One of the main conceptual strengths lies in the choice of modelling trees as individual units rather than as aggregated vegetation patches. This is particularly relevant in urban environments, where tree distribution is highly fragmented and heterogeneous (Hamberg et al., 2008). By working at the individual scale, the model captures this variability and can accommodate detailed input data from urban tree inventories, which are typically recorded at the single-tree level (Keller and Konijnendijk, 2012). This structure also makes it possible to reflect tree-specific management practices and behaviour of species-specific functional traits, improving both interpretability and practical relevance.

Given the lack of extensive urban tree related time-series (Ossola et al., 2021), full statistical approaches, as machine learning, are not suitable for urban tree modelling applications. Instead of relying only on allometric correlations, DynaTree has a mechanistic formulation, explicitly simulating biological processes, such as photosynthesis, biomass allocation, and evapotranspiration. Each component of the model is designed to reflect a cause-effect relationship based on ecological understanding, and the modular architecture enables users to activate or deactivate sub-models (e.g., for carbon sequestration, air filtration, or local climate regulation) depending on the analysis objective. Nevertheless, the allometric equations needed to estimate *DBH* from aboveground biomass, although based on literature sources, may introduce errors, particularly during the early years of growth (Blujdea et al., 2012).

The dynamic structure of DynaTree further distinguishes it from static frameworks. By simulating processes over time, it becomes possible to explore how trees grow and respond to evolving environmental conditions, including future climate scenarios. The temporal granularity is set at a daily resolution, which balances ecological realism with computational feasibility.

Moving to the limitations, the first concern biomass loss and mortality. The loss considers both natural processes (e.g., leaf fall) and human-derived interventions (e.g., branch thinning), reflecting the fact that DynaTree is calibrated using data from managed urban trees. However, biomass loss is applied using fixed coefficients, which simplifies implementation but does not capture the variability arising from environmental conditions or differences in management practices. In the same vein, the absence of tree mortality means that trees are assumed to persist throughout the simulation, which may be suitable for short-term assessments or well-managed areas, but limits the model’s realism under long-term or extreme climate scenarios. For example, simulate age-based mortality within a system dynamics framework, while (Escobedo et al., 2011) integrate species- and condition-dependent death probabilities into urban forest assessments. Introducing mortality would enhance the model’s capacity to reflect species-specific vulnerabilities and to avoid overestimating long-term ecosystem service provision. It would also capture important management implications, such as the need for replanting, handling of dead biomass, and increased costs, all of which could influence planting and policy recommendations under future climate conditions.

Another limitation lies in the simplification of processes related to stress response and phenology. For example, budburst is modelled as a temperature-dependent growing degree-days threshold, applied uniformly across species, due to the lack of detailed phenological datasets at the urban scale. Similarly, the model does not incorporate bud dormancy or chilling requirements, which could influence spring dynamics under warmer winters, a topic that remains poorly studied for many urban tree species (Cooke et al., 2012). Similarly, DynaTree does not include explicit responses to abiotic or biotic disturbances such as mechanical damage, pests, or disease, despite their known impact on tree vitality in urban settings.

The last limitation concerns the soil water balance, which is currently modelled using simplified routines for rainfall interception and infiltration. While this allows for efficient simulation across multiple years and individuals, it reduces the model’s capacity to capture fine-scale water dynamics that are especially relevant in urban settings, where soil sealing and canopy structure strongly affect water availability. More detailed representations, like those implemented in the URbanTRee model (Tams et al., 2023), account for radiation attenuation, surface types, and soil water retention in greater depth. Integrating a similar approach into this model, particularly in terms of dynamic infiltration and soil water stress responses, could substantially improve the accuracy of physiological outputs and ecosystem service estimates under variable climatic and urban conditions.

Despite these limitations, one of the strengths is that DynaTree remains as simple as possible without sacrificing essential realism, which makes it highly suitable for applied purposes. Its individual-based and modular design allows for application to other cities, provided that basic tree and climate data are available. Urban tree inventories are increasingly accessible across cities through open-data portals and remote sensing platforms (Ossola et al., 2020). This means that DynaTree can be transferred to new contexts with minimal structural adjustments, primarily by recalibrating key parameters using local datasets. The modularity will also permit to incorporate tree mortality and additional ES in future developments. For example, water flow regulation in terms of avoided run-off could be estimated by coupling canopy interception with soil infiltration dynamics, while the contribution of tree shading to local climate regulation could be derived from crown geometry and solar angle simulations. These additions would further enhance the model’s relevance for informing urban forest strategies providing a more comprehensive ES assessment.

### 4.2. Calibration and Sensitivity Analysis in Context of Existing Literature

Calibration and sensitivity analysis demonstrate that DynaTree can reproduce realistic growth trajectories and respond consistently across species. Using a multi-objective approach (NSGA-II) to calibrate both *DBH* and *CD* ensures that structural dynamics are balanced and aligned with empirical observations. Targeting the third quartile of observed values by age class helps focus the calibration on well-performing individuals, which is particularly appropriate in urban environments where stress and management practices often introduce noise into datasets. Despite the lack of individual-level time-series data, the model effectively reconstructs plausible average growth curves by aggregating cross-sectional data across age classes. While this method assumes a representative age distribution, it performs well given the available information and allows meaningful calibration even in the absence of long-term monitoring.

The sensitivity analysis confirms that DynaTree is stable under plausible parameter variation. Most parameters produce limited deviations in *DBH* and *CD* when perturbed within ±20%, indicating good structural robustness. Some parameters, such as the light extinction coefficient *k*_*2*_ and the allocation coefficient to stem *e*_*S,B*_, show a stronger influence on outputs, especially in species like *Populus nigra* and *Robinia pseudoacacia*. Comparison with literature ranges highlights some interesting points. For *k*_*2*_, our calibrated values are little lower than the typical range of 0.4–0.7 reported in forest and crop studies (Lacasa et al., 2021; Zhang et al., 2014). This difference may reflect the fact that urban trees are often isolated, so that for a given LAI more light penetrates the canopy compared to denser stands. For stem turnover, our calibrated values correspond to annual rates of 0.38–1.39%, which are slightly lower but in the same order of magnitude as those reported in mature forests (around 1.8–1.9% yr^−1^; Lewis et al., 2004; Vilanova et al., 2018). Similarly, branch turnover rates obtained here (3.7–12.8%) align reasonably well with experimental findings of highly variable branch turnover around 14% ± 19% (Lim et al., 2024). These parallels suggest that while the calibrated values are somewhat lower than those observed in unmanaged natural stands, they remain consistent in terms of magnitude, likely reflecting the fact that the model simulates ideal, well-managed urban trees rather than forest individuals.

Overall, the calibration and sensitivity analysis confirm that the model behaves coherently, responds predictably to input variation, and offers sufficient precision for exploring growth dynamics and ES supply in a realistic and controlled manner, while also producing parameter values broadly consistent with ranges documented in the literature.

### 4.3. Pilot application

The pilot application of the model in Milan serves both as a demonstration of its operational potential and as a test of its ability to reproduce plausible growth and service trajectories under contrasting climatic and planting scenarios. Overall, the results confirm that DynaTree produces realistic, species-specific dynamics that are consistent with ecological expectations and observed urban tree behaviour.

Species responses captured the expected variation in growth performance and ES supply, with the model reproducing interspecific differences under identical environmental forcing. This variability is critical for guiding species selection in urban planning, but it is also important to assess whether the magnitude of ecosystem services estimated is consistent with previous studies. For carbon sequestration, our simulations yielded average annual rates ranging from 8 kg C per tree for young individuals (5 years at planting) to 36 kg C per tree for older starting ages (20 years). These values are well aligned with the range of 6–36 kg C per tree per year reported for urban trees in Bolzano (Russo et al., 2014). For PM_10_ filtration, our estimates range from 2 g per tree per year for younger trees to 24 g per tree per year for mature individuals, corresponding to 0.4–0.8 g m^−2^ depending on species. These figures are consistent with those reported by Manes et al. (2016), who found values between 0.14 and 1.29 g m^−2^ for deciduous species in Italian cities. Finally, the estimates for heat mitigation through evapotranspiration range between 40 and 200 kWh per tree per year, depending on age and species. These are comparable with the values of 39–168 kWh per tree per year reported by McPherson et al. (1994) for trees under different exposure conditions in Chicago. Together, these comparisons suggest that the model produces realistic and ecologically plausible estimates of urban tree ecosystem services.

In terms of climate sensitivity, DynaTree simulated consistent trends across three CORDEX-based climate scenarios for Milan. However, the magnitude of variation between scenarios remained limited. This restrained divergence likely stems from the nature of the input climate data, and for the length of the projection. CORDEX models, though operate at regional scale, do not incorporate urban-specific processes, such as the urban heat island effect. As a result, the forcing data underestimate both present-day urban temperatures and the expected intensity of future warming. This limitation affects both absolute growth outcomes and the apparent impact of climate change on trees and services. More accurate urban projections would require the use of high-resolution meteorological data, ideally from urban monitoring stations or dedicated downscaling approaches that account for the complexity of the built environment. Moreover, since the current simulation spans only 25 years, the divergence between climate scenarios remains limited; longer-term projections extending to the end of the century would likely amplify these differences substantially. In this respect, this pilot application serves as a conservative estimate of what future scenarios might entail in Milan and emphasizes the need for better integration between urban climate and tree modelling.

Nonetheless, the application demonstrates DynaTree’s ability to reveal important trade-offs and nonlinear responses. For example, services like local climate regulation, which are driven by crown expansion and evapotranspiration, tend to remain stable or increase slightly under warming scenarios, even when growth rates decline. Conversely, services dependent on biomass production, such as carbon sequestration, show a stronger sensitivity to climatic constraints. These dynamics reflect the complex and service-specific responses of urban trees to environmental stress and highlight the risks of focusing on single metrics when evaluating nature-based solutions.

## 5. Conclusion

This work offers a novel contribution to urban ecological modelling by combining a mechanistic, individual-based, and dynamic approach to simulate tree growth and associated ES provision. By integrating biological processes, climate forcing, and structural dynamics, DynaTree provides a versatile framework capable of simulating long-term growth trajectories across different species and age classes. The model offers a practical balance between ecological realism and data availability, allowing simulations under various climate scenarios with limited input requirements. It performs consistently across species and outputs interpretable indicators of biomass accumulation and services such as carbon sequestration, air filtration, and local climate regulation. A key strength is its capacity to represent trees at different life stages, from saplings to centenary trees, making it applicable to planning, maintenance, and conservation alike. DynaTree structure is also aligned with the logic of the ES cascade and ecosystem accounting frameworks, ensuring that ecosystem services are calculated from simulated biophysical functioning rather than directly from land-cover proxies, which improves transparency and ecological consistency.

While the model currently simulates individual trees in isolation, future developments will aim to extend it to multiple individuals, incorporating interactions such as shading, sheltering, and belowground competition. This extension is particularly relevant for informing urban planting and management, particularly under dense tree configurations. Additional improvements will target the inclusion of further ES classes (e.g., water flow regulation, shading-based local climate regulation, habitat maintenance), mortality, and use of high-resolution urban climate data.

Overall, DynaTree provides a flexible and ecologically meaningful basis for assessing urban forestry strategies under changing climatic conditions. It supports decision-making at multiple scales, from site-level species selection and planting strategies to strategic analyses of species performance under long-term climate projections, aiding resilience planning and adaptive management. Its modularity and transparency make it a valuable tool for researchers, urban planners, and practitioners, with potential applicability to other cities where basic tree inventories and climate data are available. Combined with its conceptual structure, calibration approach, and pilot application, the model is a valuable tool for researchers, urban planners, and related practitioners working at the interface between urban ecology, climate adaptation, and ES assessment.

## Funding information

This work was supported by the National Biodiversity Future Centre (NBFC) project, funded by the European Union’s NextGenerationEU, National Recovery and Resilience Plan (NRRP), CN00000033, Concession Decree No. 1034 of 17 June 2022 adopted by the Italian Ministry of University and Research, CUP, H43C22000530001.

## Notes

### Competing Interest Statement

The authors have declared no competing interest.

