## Supplementary material for "An individual, mechanistic and dynamical model to simulate urban tree growth and ecosystem services supply under future scenarios"

### Appendix A

#### Budburst and Abscission

The parameter $a$ regulates the seasonal activation of energy allocation to leaves. Since no clear references are available for this process, we adopted a simplified formulation to approximate the natural dynamics of broadleaved deciduous trees. A very low baseline value ($a=0.01$) is assumed throughout most of the year to represent minimal allocation for maintaining buds or residual leaves. In spring, $a$ increases sharply to 0.5 to mimic budburst, and then linearly declines back to 0.01 over about 60 days, to captures the essential phenological pattern. While the shape of this curve relies on the maximum simplicity of assumptions, the timing of the onset is determined by a budburst function based on temperature accumulation (Prentice et al., 1992; Malyshev et al., 2024), specifically on growing degree-days according to the formula in Murray et al. (1989):

$$T(Budburst)=m_{1}+m_{2} exp (rC)$$

in which $T$ is the threshold of growing degree-days from which the budburst starts, $m_{1}$ is equal to -56, $m_{2}$ is equal to 602, $r$ is equal to 0.009 and $C$ is the number of chill days (temperature lower than 5 degrees Celsius) during the previous winter.

The loss coefficient $l$ varies by compartment. For leaves, $l_{L}$ includes a baseline value of 0.1% per day (Meier et al., 2006), representing a minimal temporally constant leaf loss throughout the year. For deciduous species in temperate climates, $l_{L}$ is increased in autumn, with a triangular shape to simulate seasonal abscission, reducing leaf biomass to near-zero. This adjustment was implemented manually due to the lack of quantitative references for autumnal leaf loss. The start of the abscission is set to the 1^st^ of October, the peak on the 1^st^ of November and the leaf loss is back to the value of 0.001 on the 1^st^ of December. Ideally $l_{L}$ could be linked to exogenous factors such as temperature or day-of-year, similar to the approach used for budburst. However, seasonal leaf fall has little impact on the long-term growth trajectory of the tree in this model, making this simplification appropriate for the current purpose

#### Soil characteristics

Total available water $TAW$ $\left[ mm \right]$, is the water from field capacity to wilting point that is all the water available to the plant. According to Allen et al. (1998), is calculated as

$$TAW = (\theta_{FC}-\theta_{WP})*Z_{R}$$

in which $\theta_{FC}$ is the percentage of the water in a specific soil to arrive at field capacity, $\theta_{WP}$ is the percentage of the water in a specific soil to arrive at wilting point and $Z_{R}$ is the root depth. For urban setting $Z_{R}$ is reasonably set to be 1 meter deep, while, for the specific terrain of Milan, $\theta_{FC}$ is set at 38% and $\theta_{WP}$ at 21%.

Readily available water $RAW$ $\left[ mm \right]$, is the portion of $TAW$ in which the water is free to move and so to be uptake from the plant. According to Allen et al. (1998), $RAW$ is calculated as

$$RAW = p*TAW$$

in which $p$ is the depletion factor, set to the commonly used value 0.5.

### Appendix B

#### Tree data from municipality of Milan

The city of Milan was selected as a case study for two main reasons. First, it is one of the largest urban areas in the European Mediterranean region, a zone that is expected to be particularly vulnerable to the impacts of climate change (Intergovernmental Panel on Climate Change, 2023b). Second, Milan has a well-maintained and publicly accessible inventory of urban trees in public spaces, which provides the detailed data necessary to calibrate our novel model in a context-specific and spatially explicit manner. We retrieved tree data from the Open Data portal of the Municipality of Milan (Comune di Milano, 2023). The dataset contains for each public and managed tree of the municipality information about the species, location, DBH, height, crown diameter and date of plantation. The dataset has been used for two purposes: first select species among the most common in the city (*Platanus × acerifolia*, *Populus nigra* and *Robinia pseudoacacia*), and second calibrate the model on data of these species.

#### Calibration process (in deep explanation)

The calibration made use of DBH and CD values, for the three species from data in the dataset, which are, together with height, morphological variables commonly available in urban tree inventories. Since the tree data inventory of Milan does not contain time-series on individual tree growth for the individuals of different species, neither those data are available in the literature for other relevant urban contexts, we adopted the assumption that the current snapshot of a tree species population within the city, comprising individuals of varying ages, can serve as a proxy for the average growth trajectory of that species under local urban conditions.

To smooth out gaps in the annual age distribution of trees, we group tree data for both DBH and CD into 5-year age classes, starting from 3 years old trees [3-7, 8-12, …]. For each class, the third quartile (Q3) was taken as a representative value for a healthy, well-performing tree. Calibration of the unknown parameters was then conducted using the genetic algorithm NSGA-II (Deb et al., 2002), to simultaneously minimize two objective functions, namely the mean square error between the model predictions and the third quartiles for both DBH and CD. In order to assess whether the properly calibrated model had an acceptable predictive capability, we halved the original dataset into two subsets, a calibration subset (with 50% of the data) and a validation subset (with the remaining 50%). This split ensured that the optimization process did not overfit the entire dataset and allowed for an independent evaluation of model performance. Both subsets were not generated by simple (pure) random sampling but were stratified by species and age class to preserve the overall distribution of DBH and CD values.

Calibration was conducted starting from the age class centered around 10-year-old trees, since data from younger individuals were sparse and less reliable. As a result, the initial biomass conditions at 5 years for the stem S and branches B were not fixed a priori but treated as two additional calibration parameters. Their values were constrained to fall within a biologically reasonable range (0.1 kg<B<S<5 kg).

### Appendix C

#### Climate change scenarios

The climate data were sourced from the CORDEX regional climate model ensemble (Copernicus Climate Change Service, 2019), chosen as it is the finest spatial resolution available. Specifically, we used the MPI-M-MPI-ESM-LR model downscaled with the ICTP-RegCM4-6 regional model. The values of the cell overlapping the city of Milan were extracted for the three climate scenarios selected for this pilot application. The variables that serve as inputs to our model include daily maximum, mean and minimum temperatures and precipitation. Incoming radiation was not taken directly from CORDEX but instead calculated from extraterrestrial solar radiation based on day-of-year and latitude (Allen et al., 1998). This was then attenuated using the diurnal temperature range (i.e., the difference between maximum and minimum temperature) as a proxy for cloud cover.

#### Generation of the synthetic time-series for PM10

PM_10_ concentration data were retrieved from the ARPA Lombardia monitoring station of Milano Pascal Città Studi (ARPA Lombardia, 2024), selecting 10 years of daily measurements (2011–2020). Based on these records, we generated a synthetic daily time series with equivalent statistical properties, extended to match the full simulation period required by the model. The synthetic time series has been generated with following R code:

```
# 1. Load and Convert Data into a Time Series
pm10_ts <- ts(pm10_data$PM10, frequency = 365, start = c(2011, 1))

# 2. Log-Transform the Data to Handle Skewness
log_pm10 <- log(pm10_ts + 1)  # Add 1 to avoid log(0)

# 3. Decompose the Series to Separate Components
stl_decomp <- stl(log_pm10, s.window = "periodic")

# Extract the seasonal, trend, and remainder components
seasonal <- stl_decomp$time.series[, "seasonal"]
trend <- stl_decomp$time.series[, "trend"]
remainder <- stl_decomp$time.series[, "remainder"]

# 4. Fit an ARIMA Model to the Remainder
arima_model <- auto.arima(remainder)

# 5. Simulate the Synthetic Remainder
synthetic_remainder <- simulate(arima_model, nsim = 25*365) #length(remainder)
synthetic_remainder <- ts(synthetic_remainder, frequency = 365, start = c(2025, 1))

# 5a. Create new seasonal and trend
trend<-rep(mean(trend),(365*25))
seasonal<-rep(seasonal[1:365],25)

# 6. Reconstruct the Synthetic Series
synthetic_log_series <- synthetic_remainder + seasonal +trend

# Back-transform the synthetic series to the original scale
synthetic_series <- exp(synthetic_log_series) - 1
synthetic_series <- pmax(synthetic_series, 0)  # Ensure non-negativity
```

### Tables and Figures

Table S1: mean LUE values from different species, the exact species used here were not present thus we used the median value. as an average of the value for the species used in the study

| **LUE** | **Mean** | **Min** | **Q1** | **Median** | **Q3** | **Max** | **Dev** |
| --- | --- | --- | --- | --- | --- | --- | --- |
| 1 | 0.06023548 | 0.012 | 0.023 | 0.063 | 0.08043078 | 0.24 | 0.04947834 |

Table S2: SLA value from TryDB

| **SpeciesName** | **Mean** | **Min** | **Q1** | **Median** | **Q3** | **Max** | **Dev** |
| --- | --- | --- | --- | --- | --- | --- | --- |
| Platanus x acerifolia | 17.380 | 7.576 | 14.500 | 15.038 | 23.277 | 23.280 | 5.389 |
| Populus nigra | 13.811 | 10.070 | 10.600 | 13.961 | 16.630 | 19.000 | 3.299 |
| Robinia pseudoacacia | 24.361 | 9.705 | 18.901 | 21.383 | 26.400 | 56.000 | 10.827 |

Table S3: allometric equation parameters used for different species. As a rule, we first selected the species-related parameters in urban environment to obtain the best available allometric equation. In cases where these are not available, we parameterize the equation using data for the same species living in forested habitats with similar climates.

| **Species** | **a** | **b** | **c** | **d** | **Reference** |
| --- | --- | --- | --- | --- | --- |
| Platanus x acerifolia | 0.0295 | 2.67 | 3.742 | 0.375 | [1] McPherson, 2016, Urban tree database and allometric equations, https://doi.org/10.2737/PSW-GTR-253 |
| Populus nigra | 0.1177 | 2.26 | 2.578 | 0.442 | [4] Fortier, 2017, Allometric Equations for Estimating Compartment Biomass and Stem Volume in Mature Hybrid Poplars: General or Site-Specific?, 10.3390/f8090309 |
| Robinia pseudoacacia | 0.1075 | 2.39 | 2.543 | 0.418 | [5] Annighöfer, 2012, Biomass functions for the two alien tree species Prunus serotina Ehrh. and Robinia pseudoacacia L. in floodplain forests of Northern Italy, 10.1007/s10342-012-0629-2 |

Figure S1: Influence of calibrated parameters k₁ and k₂ on morphological traits. (a) Relationship between total biomass and Crown Projected Area (CPA), as governed by k₁. (b) Relationship between Leaf Area Index (LAI) and fraction of light intercepted, as shaped by k₂. Species-specific differences reflect varying growth strategies and light-use efficiencies between emprical parameters and tree variables.

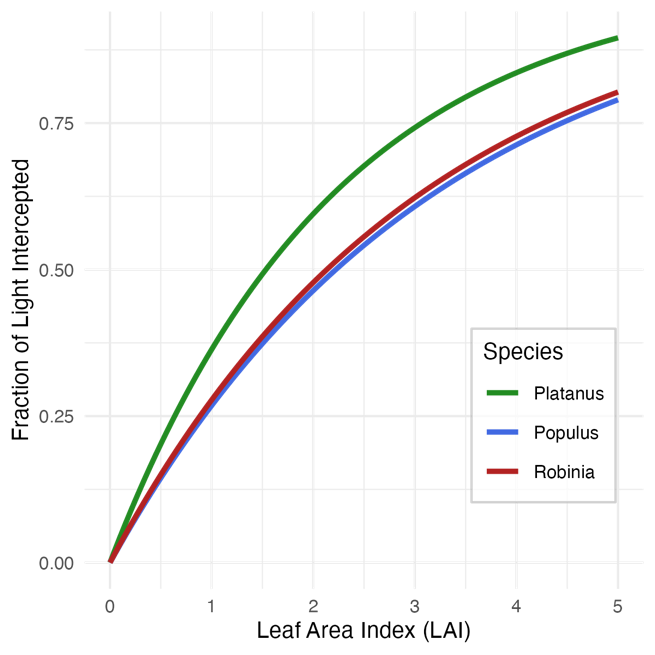

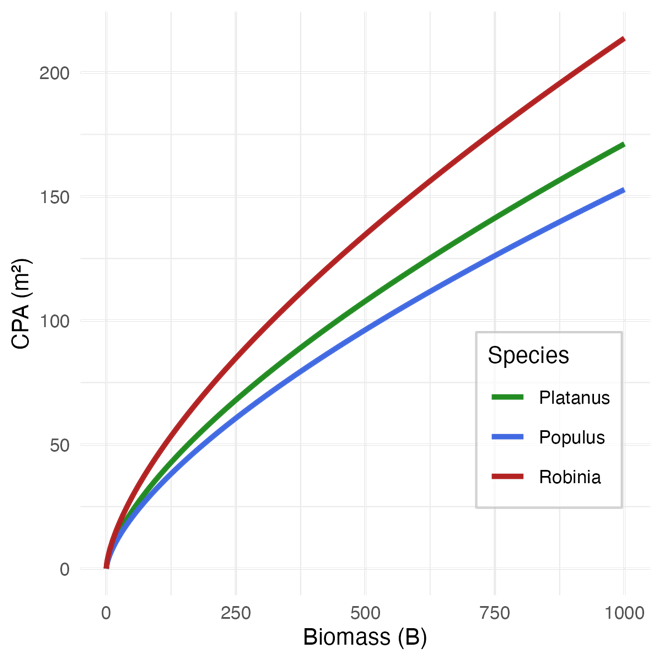
